# Buzzing Frequency Influences Pollen Release in Buzz-Pollinated Poricidal Anthers

**DOI:** 10.64898/2026.06.17.732968

**Authors:** Mitchell Alvord, Braden Cote, Sarah Morris, Mark Jankauski

## Abstract

Buzz pollination is an important behavior in which bees use vibrations to extract pollen from poricidal anthers. However, the extent to which vibration frequency influences pollen release remains unclear. Here, we quantified pollen expulsion from *Solanum sisymbriifolium* anthers subjected to harmonic excitation over a broad frequency range encompassing the anther’s first natural frequency. We excited anthers to expel pollen and measured anther kinematics and pollen release using high-speed videography. Particle tracking enabled continuous estimation of pollen release throughout each buzzing event, allowing both initial pollen flux and total pollen released to be quantified. Pollen release depended strongly on excitation frequency. Initial pollen flux, total pollen release, and anther kinematics peaked when excitation frequency approached the anther’s natural frequency. Anther tip velocity amplitude exhibited the strongest correlation with total pollen release (*r* = 0.755) and initial pollen flux (*r* = 0.898). Experimental observations were compared with nonlinear and linear statistical models of pollen release. While both models captured trends in normalized pollen flux, they overpredicted total pollen release, suggesting that adhesive interactions play important roles during extended buzzing events. These findings demonstrate that anther structural dynamics influence pollen release and suggest that vibration amplification may improve the efficiency of buzz pollination.

## 1 Introduction

Buzz pollination is a plant-pollinator interaction in which bees use mechanically generated vibrations to release pollen from flowers with poricidal anthers [1–8]. Poricidal anthers are hollow, tube-like structures that retain pollen internally and permit its dispersal only through small apical openings [9– 11]. Approximately 10% of flowering plant species possess these specialized anthers [12, 13]. During buzz pollination, a bee grasps the anther and rapidly activates its indirect flight muscles, producing high-frequency vibrations that are transmitted to the anther [14, 15]. These vibrations mobilize pollen within the anther and eject it through the flower’s apical pores, where it adheres to the bee and is subsequently transported to other flowers. Buzz pollination is performed by a diverse array of bee species and is required for the pollination of more than 20,000 flowering plant species worldwide [16]. Many economically important crops, including tomatoes, blueberries, and kiwifruit, rely on this behavior for successful pollination [17]. Beyond its agricultural significance, buzz pollination plays an ecological role by supporting the reproduction of wild plant communities and the pollinators that depend on them [18].

The effectiveness of buzz pollination depends on the mechanical interaction between the bee and the poricidal anther [19]. Floral buzzing occurs in repeated bursts as a bee grasps the anther and rapidly activates its indirect flight muscles [20]. These bouts of buzzing generate large oscillatory forces that excite the flexible anther and induce the deformations responsible for pollen release. The forces produced during buzzing may be considerable; for example, forces generated during defensive buzzing, a non-flight behavior driven by the same flight muscles used in floral buzzing, have been measured at up to 80 times the body weight of large carpenter bees [21]. Buzzing frequencies vary among pollinator species but are typically reported in the range of 100–400 Hz [22]. Such vigorous vibration production, typified by large forces applied at high frequency, is energetically expensive. Estimates of the mass-specific energetic cost of floral buzzing in the bumblebee Bombus terrestris approach 300 W/kg [23], comparable to reported energetic costs of flight. This suggests that buzz pollination may represent a substantial energetic investment for bees that routinely forage from flowers with poricidal anthers. However, the effectiveness of this energetic investment ultimately depends on how the applied forces are transmitted through the flower and converted into anther deformation.

The resulting anther deformation depends not only on the magnitude of the applied forces but also on the structural dynamics of the anther itself. A key dynamic property is the natural frequency, defined as the inherent frequency at which the anther tends to vibrate when disturbed [24]. Anther natural frequencies are governed by their morphology, material properties, and turgor pressure [25, 26]. When the excitation frequency approaches the natural frequency, resonant amplification can occur, resulting in comparatively large deformations even under relatively small oscillatory forces. Previous studies have reported natural frequencies of *Solanum sisymbriifolium* in the range of about 500-750 Hz [26], generally outside the frequencies observed during buzz pollination. However, these estimates neglect the mass of the pollinating bee. Computational studies suggest that bee mass can substantially reduce anther natural frequencies, in some cases shifting them into the range of floral buzzing frequencies [25]. Consequently, the extent to which resonance contributes to buzz pollination, and whether such excitation enhances pollen release, remains unclear.

Several studies have investigated how vibration influences pollen release from poricidal anthers. In multiple species within the *Solanum* genus, pollen dispensing during repeated bouts of artificial buzzing follows an exponential decay pattern, with progressively smaller quantities of pollen released during successive vibrations [27]. Similar behavior has been observed under natural pollination conditions; for example, *Solanum rostratum* flowers visited by the bumblebee *Bombus terrestris* release only approximately 2–4% of their pollen during each second of buzzing [28]. Together, these observations suggest that pollen dispensing is a gradual process that unfolds over multiple buzzing events. Several factors have been proposed to influence the rate and extent of this dispensing process. Longer buzzing durations and higher vibration velocity amplitudes generally increase pollen removal [29–31], while floral morphology also plays an important role. For example, flowers with fused anthers tend to release more pollen than species with loosely arranged anthers [32], and smaller flowers often exhibit faster pollen dispensing schedules than larger flowers [27]. In contrast, several studies have reported that buzzing frequency has little or no direct effect on pollen release [29, 30, 33]. This conclusion is surprising because vibration frequency is a primary determinant of anther dynamic response and therefore strongly influences the vibration amplitudes experienced by the flower. If pollen release depends on anther kinematics, as previous work suggests, then excitation frequency may influence pollen release indirectly through its effect on anther dynamics. Consequently, the extent to which buzzing frequency affects pollen release, particularly when excitation approaches the natural frequency of the anther, remains unresolved.

The goal of this study is to determine whether buzzing frequency influences pollen expulsion from poricidal anthers. To this end, we developed a novel experiment to simultaneously measure anther deformation and pollen release in artificially excited *Solanum sisymbriifolium* anthers. Anthers were excited using an electrodynamic shaker over a broad frequency range encompassing their first natural frequency, while high-speed videography was used to quantify both anther kinematics and pollen expulsion. The use of high-speed videography enables pollen release to be measured nearly continuously throughout a buzzing event, in contrast to previous studies that quantified expelled pollen only at discrete intervals. Pollen release was characterized using two metrics: (1) the initial pollen flux measured at the onset of buzzing and (2) the total pollen released during a one-second buzzing event. These metrics were evaluated as functions of vibration frequency and anther tip displacement, velocity, and acceleration amplitudes. Experimental observations were compared with a statistical pollen release model adapted from a previously published biophysical framework. Finally, we interpret our findings within the broader context of buzz pollination, hypothesizing that anther structural dynamics play a fundamental role in pollen release and discussing how existing biophysical models may be refined to better predict pollen ejection.

## 2 Methods

### 2.1 Study Species and Preparation

We grew *Solanum sisymbriifolium* plants in the Plant Growth Center at Montana State University. Once they began flowering, we cut the fresh blooms at the pedicel and immediately inserted them into centrifugal tubes filled with water to prevent wilting and maintain turgor. We transported the flowers to the laboratory and tested them on the same day as harvest. In total, we tested *n* = 110 anthers.

The mounting and excitation configuration is shown in Fig. 1A. We prepared stamens by excising them from the flower. We glued the base of each anther to a 3D-printed cube, with the stamen filament extending into a central hole within the cube. We then mounted the cube in an inverted orientation within a clamp attached to an optical breadboard. To excite the anther, we used a 31-N (7-lbf) electrodynamic shaker (K2007, The Modal Shop, Cincinnati, OH, USA) mounted on a translation stage and powered by an external amplifier (2100E23-100, The Modal Shop, Cincinnati, OH, USA). We transmitted vibrations to the anther through a fine gage insect pin attached to the shaker armature. Using the translation stage, we brought the pin into contact with the anther wall, secured the connection with a small amount of super glue (Gorilla Super Glue, The Gorilla Glue Company, Cincinnati, OH, USA), and allowed it to cure for five minutes before testing. During experiments, we enclosed the assembly within a shield featuring two translucent Plexiglas walls to minimize the influence of room air currents on pollen motion and release.

**Figure 1:**
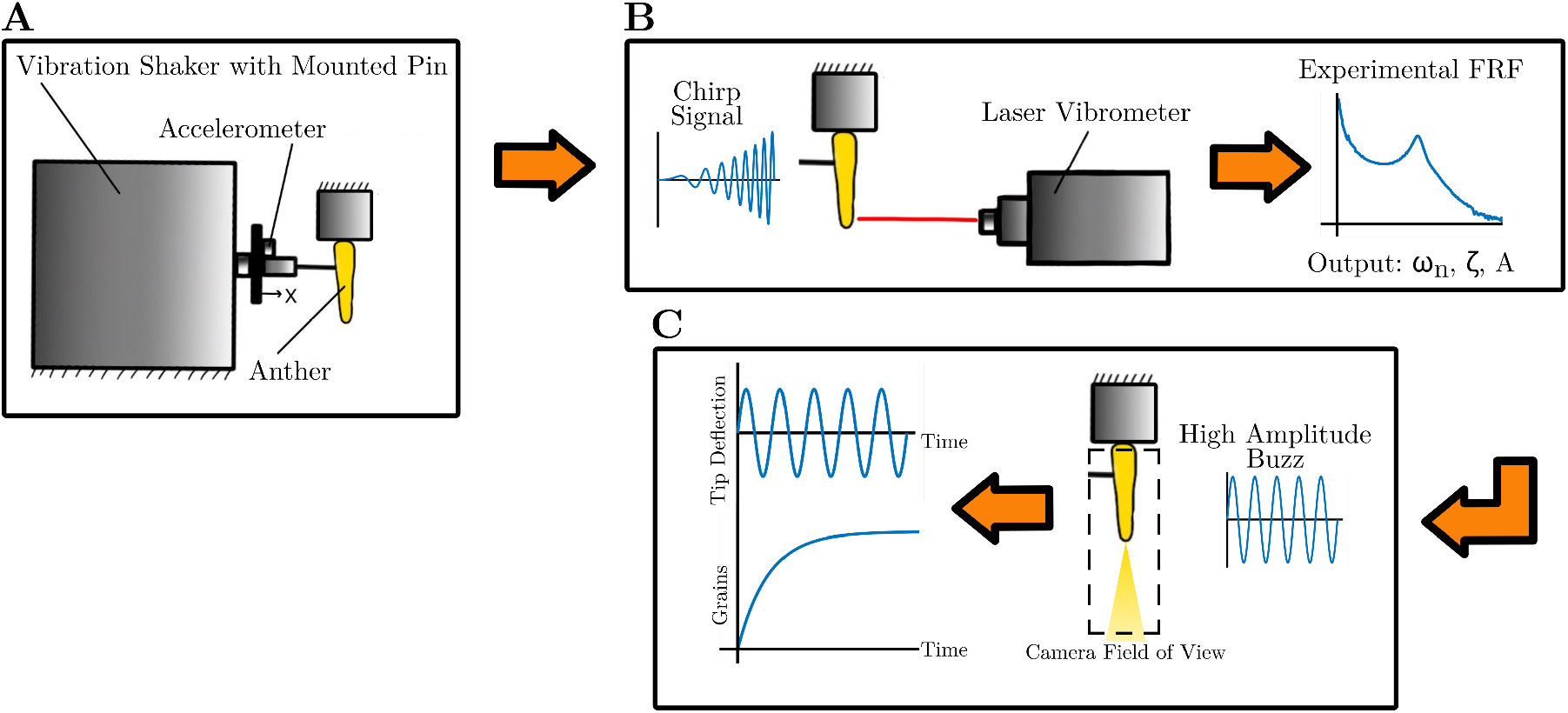
Experimental workflow and representative outputs. (A) Experimental setup used to apply prescribed vibrations to mounted anthers. An accelerometer measured the base acceleration of the shaker armature. (B) Experimental modal analysis procedure, in which a chirp excitation and laser vibrometer measurements were used to obtain the frequency response function (FRF) and identify the natural frequency *ω*_*n*_, damping ratio *ζ*, and gain constant *A*. (C) High-amplitude vibration experiments used to quantify pollen release. Tip displacement and cumulative pollen release were determined during post-processing using motion-tracking and particle-tracking techniques, respectively.

### 2.2 Experimental Techniques

#### 2.2.1 Experimental Modal Analysis

We performed low amplitude experimental modal analysis (EMA) to characterize the dynamic properties of individual anthers prior to pollen expulsion testing (Fig. 1B). Specifically, we measured the frequency response function (FRF) relating input acceleration at the excitation point to output velocity at the anther tip and used the resulting FRF to estimate the anther’s first natural frequency and damping ratio. We restricted vibration amplitudes during EMA to ensure that no pollen was released.

To measure the FRF, we used a dynamic signal analyzer, DSA, (Abacus 901, Data Physics, Riverside, CA, USA) to excite the shaker and record data. We supplied the shaker with a periodic chirp spanning a frequency range of 0 to 2000 Hz with a duration of 640 milliseconds. Data were collected over 10 excitation periods and subsequently averaged. The input acceleration was measured using an accelerometer (352A21, PCB Piezotronics, Depew, NY, USA) with a sensitivity of 10.2 mV/g. Anther tip velocity was measured using a laser vibrometer (VibroGo, Polytec, Hudson, MA, USA) with a sensitivity of 25 mm/s-V. The anther velocity and reference acceleration were converted from the time domain to the frequency domain via the DSA’s Fast Fourier Transform algorithm, and their ratio resulted in the anther FRF.

We determined the anther’s natural frequency *ω*_*n*_ and damping ratio *ζ* by modeling the anther and shaker system as a base excited, single-degree-of-freedom system and curve fitting the magnitude of the experimentally measured FRF. Parameters of the analytical FRF were estimated using a nonlinear least squares regression in Matlab’s R2024a curve fitting toolbox. The magnitude of the analytical frequency response relating input acceleration *A*_*ref*_ to anther tip velocity *V*_*tip*_ is given by:

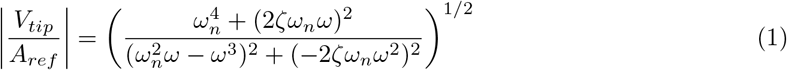

Knowledge of the anther’s natural frequency *ω*_*n*_ enables us to represent output measurements, including anther kinematics and pollen release metrics, as functions of the ratio of excitation frequency to natural frequency rather than absolute excitation frequency.

#### 2.2.2 High Amplitude Pollen Expulsion

Following low-amplitude EMA, we performed high-amplitude vibration experiments to excite the anther with sufficient energy to release pollen (Fig. 1C). These tests were designed to determine whether pollen release depends on excitation frequency and its proximity to the anther’s natural frequency. We excited each anther harmonically for a 1-s burst at a single excitation frequency. Excitation frequencies ranged from 300 to 1300 Hz in 100-Hz increments. We tested 10 anthers at each frequency, yielding a total sample size of *n* = 110. We controlled the excitation velocity to approximately 40 mm/s (40.02 ± 0.63 mm/s, mean ± SD) using a function generator (SDG1025, Siglent, Shenzhen, China) and shaker amplifier.

We recorded both pollen release and anther tip kinematics using a high-speed camera (Chronos 2.1, CR21-HD, Kron Technologies, Burnaby, BC, Canada) and a macro lens (Rokinon 100mm F2.8 Full Frame Macro Lens, Rokinon, New York, USA). All video recordings were obtained at 5406 frames/s with a 178 microsecond exposure time, f/18 aperture and a camera resolution of 480 × 800 pixels. Recorded videos were saved as grayscale 8 bit videos using the H.264 MPEG-4 encoding save format on the camera. We positioned the camera at a fixed distance from the anther such that the entire anther, including its glued base, remained within the field of view while providing sufficient space below the anther tip to capture expelled pollen. We illuminated the specimen using two studio lights (SL-200W, Godox, Shenzhen, China). We aligned the focal plane with the longitudinal axis of the anther to maximize image sharpness. We simultaneously recorded the shaker input using the same accelerometer employed during EMA testing (352A21, PCB Piezotronics, Depew, NY, USA).

### 2.3 Data Post Processing

#### 2.3.1 Video Pre-Processing

We pre-processed videos from the high-amplitude pollen expulsion experiments by extracting individual image frames for subsequent kinematic and pollen-tracking analyses. We saved frames in TIFF format to preserve image quality and avoid compression artifacts. We converted tracking outputs from pixels to SI units using a calibration factor of 0.0167 mm/pixel determined from a ruler positioned at the anther location prior to testing, which corresponded to a field of view window of approximately 8 mm × 13.4 mm over all tests. Because the anther position and pin location did not vary greatly over trials, the calibration factor was held constant for all tests. The resulting calibrated image sequences served as the basis for the quantification of both anther tip kinematics and pollen expulsion, as described in the following sections.

#### 2.3.2 Anther Tip Kinematics

We quantified anther tip kinematics to (1) determine whether excitation near the natural frequency elicited larger amplitude responses and (2) identify relationships between anther motion and pollen expulsion.

We used DeepLabCut, an open-source markerless motion-tracking framework, to track the position of the anther tip in each video frame [34]. We trained the network using ten videos of anthers excited over a frequency range of 300–1200 Hz, one at each 100-Hz increment. We manually annotated a subset of frames from these videos and used them to train the model. The trained network provided the anther tip position as a function of time for each experiment.

We analyzed the resulting trajectories in MATLAB R2024a using the Curve Fitting Toolbox. Specifically, we fit the tracked tip motion to a single-term harmonic function and extracted the oscillation frequency and displacement amplitude from the fitted model. Across all excitation frequencies, the mean coefficient of determination was *r*^2^ = 0.929 ± 0.098, suggesting that the harmonic model accurately described the measured motion. We subsequently computed velocity and acceleration amplitudes by multiplying the displacement amplitude by the angular frequency, *ω*, and angular frequency squared, *ω*^2^, respectively.

#### 2.3.3 Pollen Particle Tracking

We estimated pollen release as a function of time using particle-tracking methods. A graphical overview to the approach used to track particles is shown in Fig. 2. From these measurements, we estimated both the initial pollen flux at the onset of excitation and the total pollen expelled during the 1-s excitation burst. Because clumping or overlapping pollen grains may be interpreted as a single particle via tracking methods, we perform area-based calculations and estimate pollen grain count based on an area projection of a single pollen grain. The projected area of one pollen grain was estimated using SEM images and found to be 697.9 *μm*^2^ . Although this image-based approach may introduce greater uncertainty than direct particle-counting methods, such as a particle counter or hemocytometer, it enabled us to obtain time-resolved estimates of pollen release that were necessary for quantifying initial pollen flux. Further, because all videos were acquired and analyzed using the same imaging and tracking parameters, any systematic errors associated with the image-based counting procedure are expected to be similar across trials. Consequently, this approach remains well suited for comparing relative differences in pollen release metrics among experimental conditions.

**Figure 2:**
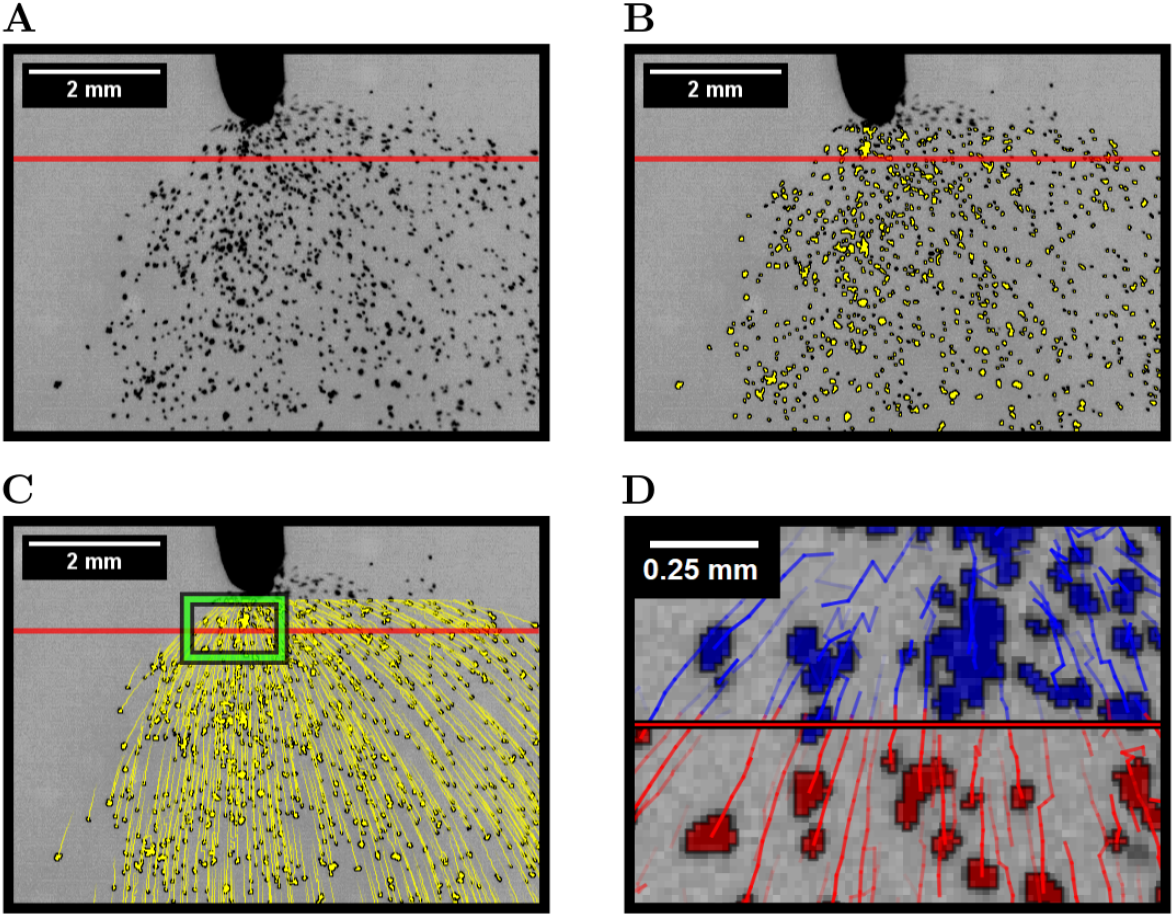
Example of processing for tracking particle grains and clumps. Images were inverted and contrast enhanced to improve visibility in the published figure. The red line in each image represents the virtual boundary that particles were counted over, located 0.5 mm from the anther tip. (A) represents the original output frame from the recorded video. (B) threshold detection done to differentiate particles from background. (C) The trajectories overlaid onto the detected particles. The tails of the tracks are backward in time and represent the particles centroid position for the last ten frames. (D) Zoomed region representing the green inset shown in panel C. Shows the trajectories crossing the virtual boundary. The tails of the particles represent the previous ten frames of the tracked particles. Blue particles are spots that are before the virtual boundary and red particles have crossed the boundary.

We analyzed video frames in the open-source image-processing software FIJI [35]. We tracked pollen particles using the native TrackMate plugin and its Linear Assignment Problem (LAP) tracking algorithm [36, 37]. Prior to tracking, we cropped each video to retain only the region immediately below the anther tip where pollen release occurred. We detected pollen grains using TrackMate’s threshold-based detector, which identifies particles based on their grayscale intensity relative to the background. We set the detection threshold to a pixel intensity of 45 and held this value constant across all experiments. For each detected particle, the algorithm recorded both its position and projected area in every frame.

The LAP tracker employs a global optimization approach that minimizes the total linking cost across detections in successive frames, with spatial distance serving as the primary cost metric. We set the maximum frame-to-frame linking distance to 5 pixels for all videos. We disabled gap closing, track splitting, and track merging for all analyses. TrackMate assigned a unique track ID to each detected particle and linked its trajectory across successive frames. We analyzed the resulting trajectories in MATLAB to quantify pollen release. Specifically, we counted particles as they crossed a virtual boundary positioned approximately 0.5 mm below the anther tip. The cumulative number of particles crossing this boundary was subsequently used to estimate pollen release as a function of time.

The virtual-boundary analysis yielded the cumulative pollen expelled from the anther as a function of time, *P* (*t*). We modeled this quantity using an exponential function. Previous studies have successfully used exponential models to describe pollen release from poricidal anthers over discrete buzzing intervals [27]. Accordingly, we expressed cumulative pollen release as

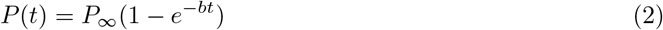

where *P*_*∞*_ is the total amount of pollen expelled at the conclusion of the test and *b* is a parameter that characterizes the rate of pollen release. The mean coefficient of determination across all fitted trajectories was *r*^2^ = 0.984 ± 0.013 (mean ± SD), indicating that the exponential model accurately described the observed release dynamics. To estimate the initial pollen flux for each test, we evaluated the time derivative of *P* (*t*) at *t* = 0. The resulting initial flux 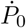 is

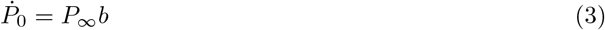

### 2.4 Modeling

#### 2.4.1 Nonlinear Model of Pollen Release

We adopted a statistical model of pollen release to quantify the relationship between anther tip kinematics and the resulting pollen release metrics. While available higher-fidelity approaches, such as billiards-based models [38, 39] and discrete element methods [40, 41], can provide detailed predictions of individual pollen trajectories, they are primarily computational and do not yield closed-form expressions relating anther motion to pollen release.. In contrast, the statistical framework used here provides analytical expressions that relate pollen flux to measurable kinematic quantities. This allows us to compare model predictions directly with experimental observations and evaluate the extent to which simple kinematic relationships explain pollen release dynamics.

We modified the biophysical model of pollen release initially formulated by Buchmann and Hurley [13]. The model treats pollen grains as an ideal granular gas, neglecting interactions between individual grains. The pollen chamber is represented as a rectangular prism of volume V containing an aperture of area *A*^*′*^ that represents the apical pore area through which pollen escapes. The chamber is subjected to a uniform prescribed translational velocity, *V*, that represents the vibration generated during buzz pollination. In its original formulation, the model assumes that the entire pollen chamber undergoes uniform translational motion. Consequently, it does not account for the spatially varying deformations observed in vibrating anthers. To facilitate comparison with our experiments, we prescribed the chamber velocity using the measured anther tip velocity, *V*_*tip*_. Pollen grains gain energy through collisions with the vibrating chamber walls, each having surface area *A*_*W all*_. The model characterizes these collisions using a coefficient of restitution, *e*, defined as the ratio of a particle’s velocity after collision to its velocity before impact.

Two coupled, nonlinear differential equations describing the mean particle energy density *ε* and particle density *η* are derived using an energy-balance formulation. In non-dimensional forms, the energy density *ε*^*′*^, particle density *η*^*′*^ and time scale *τ* are

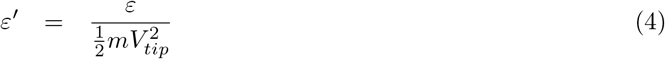

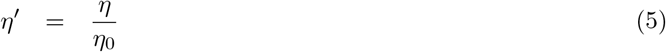

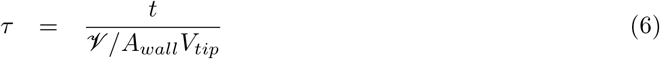

where *m* is the mass of a pollen grain and *η*_0_ is the initial particle density. The non-dimensional differential equations governing *ε*^*′*^ and *η*^*′*^ are

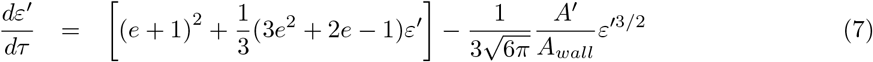

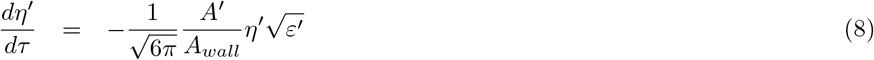

Above, the first equation represents the non-dimensional change in pollen energy density with respect to dimensionless time. The second equation represents the change in non-dimensional particle density with respect to dimensionless time.

#### 2.4.2 Approximate Linear Model of Pollen Release

Here, we derive a closed-form analytical approximation to predict pollen release metrics. Previous applications of this framework have relied on numerical integration of Eqs. 7-8 owing to their nonlinear structure. The analytical approximation derived here provides direct insight into how pollen flux depends on anther geometry and vibration kinematics, while enabling straightforward comparison with experimental observations. In the Results, we compare predictions of the approximate analytical model with those of the full nonlinear model to assess the validity of the simplifying assumptions introduced below.

Consider the non-dimensional change in pollen energy with respect to dimensionless time described by Eq. 7. The first term is a constant and represents the rate at which energy is injected to the system due to wall collisions. The second term represents an energy level correction factor that depends on the current energy per particle *ϵ*, where *ϵ* must be positive. This term is negative when 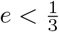 and positive when 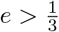 . The third term represents the reduction in particle energy as pollen grains escape from the anther and is always negative. Prior models show that when *e* is large, the rates at which pollen leaves the anther are considerably higher than what is observed in natural systems. Thus, we restrict our model to low coefficients of restitution between 0 ≤ *e* ≤ 0.2. For 0 ≤ *e* ≤ 0.2, the coefficient multiplying the second term in Eq. 7 remains *O*(10^*-*1^), whereas the prefactor associated with the third term is proportional to *A*^*′*^*/A*_*wall*_. For biologically realistic anthers, *A*^*′*^*/A*_*wall*_ is small, typically ranging from about 0.5% to 5% in some *Solanum* species [27]. Consequently, the characteristic timescale governing the evolution of particle energy is expected to be substantially shorter than that governing particle loss. We therefore assume that *ϵ*^*′*^ rapidly approaches a steady value while *η*^*′*^ evolves more slowly. Thus, we assume that the third term of Eq. 7 is negligible and that pollen energy density, 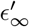, can be modeled as a constant satisfying

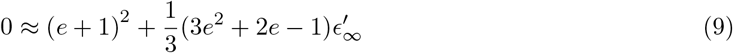

Solving the above for 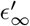 yields an estimate for constant pollen energy density 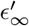 as

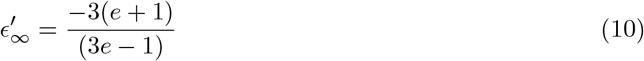

By treating the pollen energy density *ϵ*^*′*^ as a constant such that 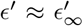, Eq. 8 becomes linear in *η*^*′*^ and is now represented as

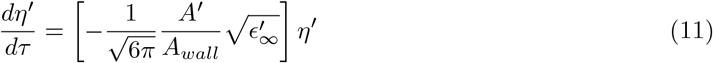

The solution for this linear ordinary differential equation governing *η*^*′*^ is then readily obtained via separation of variables as

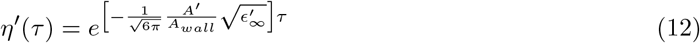

#### 2.4.3 Implementation of Pollen Release Models

We compare pollen release metrics predicted by the nonlinear and linear models to one another and to experimental measurements. Unless otherwise noted, both models employ identical parameter values to ensure a consistent comparison. Following the assumptions introduced in Section 2.4.2, we consider coefficients of restitution of *e* = 0 and *e* = 0.2. We consider an aperture-to-wall area ratio of *A*^*′*^*/A*_*wall*_ = 0.01. The pollen chamber is approximated as a rectangular prism with width *w* = 1.15 mm and length *L* = 8.6 mm based on morphometric measurements of *Solanum sisymbriifolium* reported in [26]. This approximation treats the four pollen cavities identified in [26] as a single equivalent pollen chamber. The resulting chamber volume is V = *Lw*^2^ and the wall area is *A*_*wall*_ = *Lw*. Both models are evaluated over vibration velocity amplitudes of ∈ *V*_*tip*_ [0, 1000] mm/s, consistent with the experimentally measured anther tip velocities.

For the nonlinear model, Eqs. 7-8 were solved numerically in MATLAB R2024a using the built-in ODE45 solver. The initial conditions were specified as *ε*^*′*^(0) = 0 and *η*^*′*^(0) = 1, corresponding to particles initially at rest and a fully populated pollen chamber. Simulations were performed over the dimensionless time interval *τ* ∈ [0, 3000] using adaptive time stepping. The resulting solution *η*^*′*^(*τ* ) was subsequently dimensionalized in time to obtain the particle density as a function of physical time, *η*^*′*^(*t*). Differentiation of *η*^*′*^(*t*) yielded the predicted pollen flux, *dη*^*′*^(*t*)*/dt*. Because particles initially gain energy through collisions with the vibrating chamber walls, the maximum flux does not occur immediately at *t* = 0. We therefore identify the maximum value of *dη*^*′*^(*t*)*/dt* and report this quantity as the predicted initial pollen flux. To quantify pollen release over a finite buzzing event, we additionally evaluate *η*^*′*^(*t*) at *t* = 1 s.

For the linear model, we re-dimensionalize Eq. 12 in time to obtain

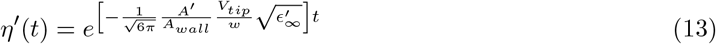

The particle density remaining after a 1-second buzzing event is therefore

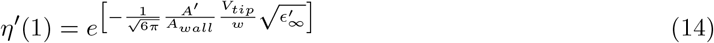

Differentiating with respect to time yields the change in particle density with respect time

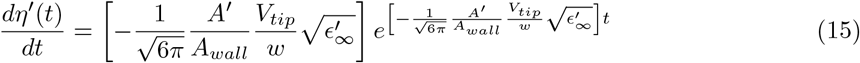

Because the exponential term decreases monotonically with time, the maximum change in particle density occurs at *t* = 0. The corresponding maximum rate of change of particle density is therefore

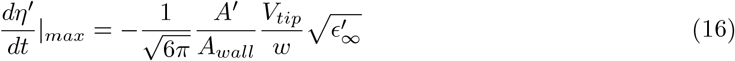

Thus, both the particle density remaining after a 1-second buzzing event and the maximum rate of change of particle density with respect to time can be determined directly from the model parameters without numerical integration.

Finally, we normalize our experimental results to enable comparison against the models. Recall that from the videographic methods, we are estimating total pollen particles ejected *P* (*t*) rather than dimensionless particle density *η*^*′*^(*t*). The relationship between *P* (*t*) and *η*^*′*^(*t*) is

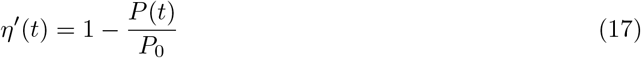

where *P*_0_ denotes the initial number of pollen grains contained within the anther prior to buzzing. Direct estimation of *P*_0_ as not possible in the present experiments because it would require quantifying both the grains ejected during buzzing and those remaining within the anther afterward. To facilitate comparison among trials and with the model predictions, we therefore normalize pollen counts by the maximum number of grains expelled in any trial, *P*_*max*_ = 49,799 grains. Despite its limitations, this approach provides a common experimental scale for pollen release, with *P*_*max*_ serving as a proxy normalization factor rather than a direct estimate of the true initial pollen count.

## 3 Results

### 3.1 Optimal frequency for pollen release

Pollen release depended strongly on buzzing frequency and its proximity to the anther natural frequency (Fig. 3 A,B). Across all trials, both the total amount of pollen released and the initial pollen flux increased with frequency ratio, reached a maximum near the anther natural frequency, and subsequently declined at higher frequency ratios. The mean natural frequency of the anthers (*n* = 110) was 876 ± 117 Hz, with a mean damping ratio of 0.075 ± 0.01. The total number of pollen grains released averaged 17,109 ± 11,409 grains and ranged from 151 to 49,799 grains across all trials. The upper bound of pollen grains released is similar to the total number of pollen grains in *Solanum rostratum* anthers [27]. Initial pollen flux exhibited even stronger frequency dependence, reaching a maximum of 184,805 grains/s at a frequency ratio of approximately 0.92 and decreasing substantially at both lower and higher frequency ratios. The tendency for pollen release metrics to peak near the natural frequency suggests that the underlying mechanical response of the anther plays an important role in governing pollen ejection.

**Figure 3:**
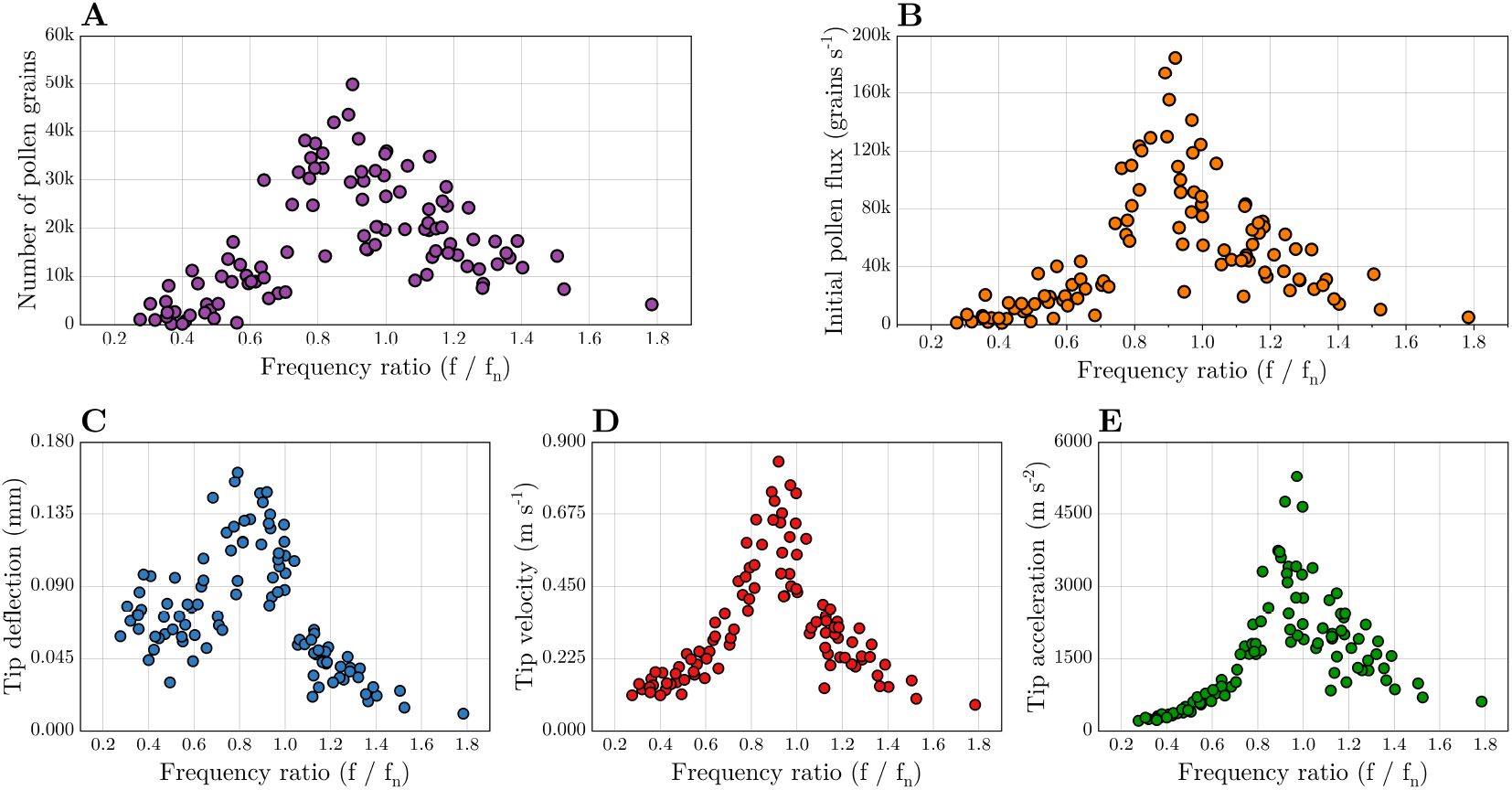
Pollen release and anther dynamics plotted against the frequency ratio. The frequency ratio is defined as the applied buzzing frequency divided by the anther’s natural frequency. (A) The total number of pollen grains expelled over the 1-second buzzing duration vs normalized frequency. (B) Initial pollen flux vs normalized frequency. (C,D,E) Tip displacement, velocity and acceleration amplitudes vs normalized frequency.

### 3.2 Resonant amplification of anther vibration

Anther tip kinematics exhibited pronounced resonant amplification when excited near the natural frequency (Fig. 3 C-E). Tip displacement, velocity, and acceleration amplitudes all increased with frequency ratio and reached maxima near *f/f*_*n*_ ≈ 0.9. This maxima occurs lower than the expected frequency ratio of *f/f*_*n*_ = 1, likely because the loss of turgor pressure between the EMA and pollen expulsion trial reduces the natural frequency of the anther. The largest measured tip displacement, velocity, and acceleration amplitudes were 0.161 mm, 0.84 m/s, and 5,285 m/s^2^, respectively. The amplification was particularly evident in the tip velocity response, which increased approximately five-fold relative to responses observed at low and high frequency ratios. Relative to the constant shaker input velocity amplitude of 40 mm/s, the maximum measured tip velocity corresponded to an amplification factor of approximately 21. Together, these results demonstrate that small changes in buzzing frequency near the natural frequency can substantially alter the kinematic response of the anther.

### 3.3 Pollen release characteristics versus anther tip kinematics

Pollen release metrics increased with increasing anther tip kinematics (Fig. 4). Larger tip displacement, velocity, and acceleration amplitudes generally resulted in greater total pollen release and higher initial pollen flux. The largest pollen release metrics were consistently associated with the largest vibrational amplitudes. Variability in pollen release also increased with kinematic amplitude, particularly for total pollen released. At the highest tip velocities, total pollen release appeared to approach a plateau, suggesting that pollen availability may have limited further increases in pollen expulsion for some anthers.

**Figure 4:**
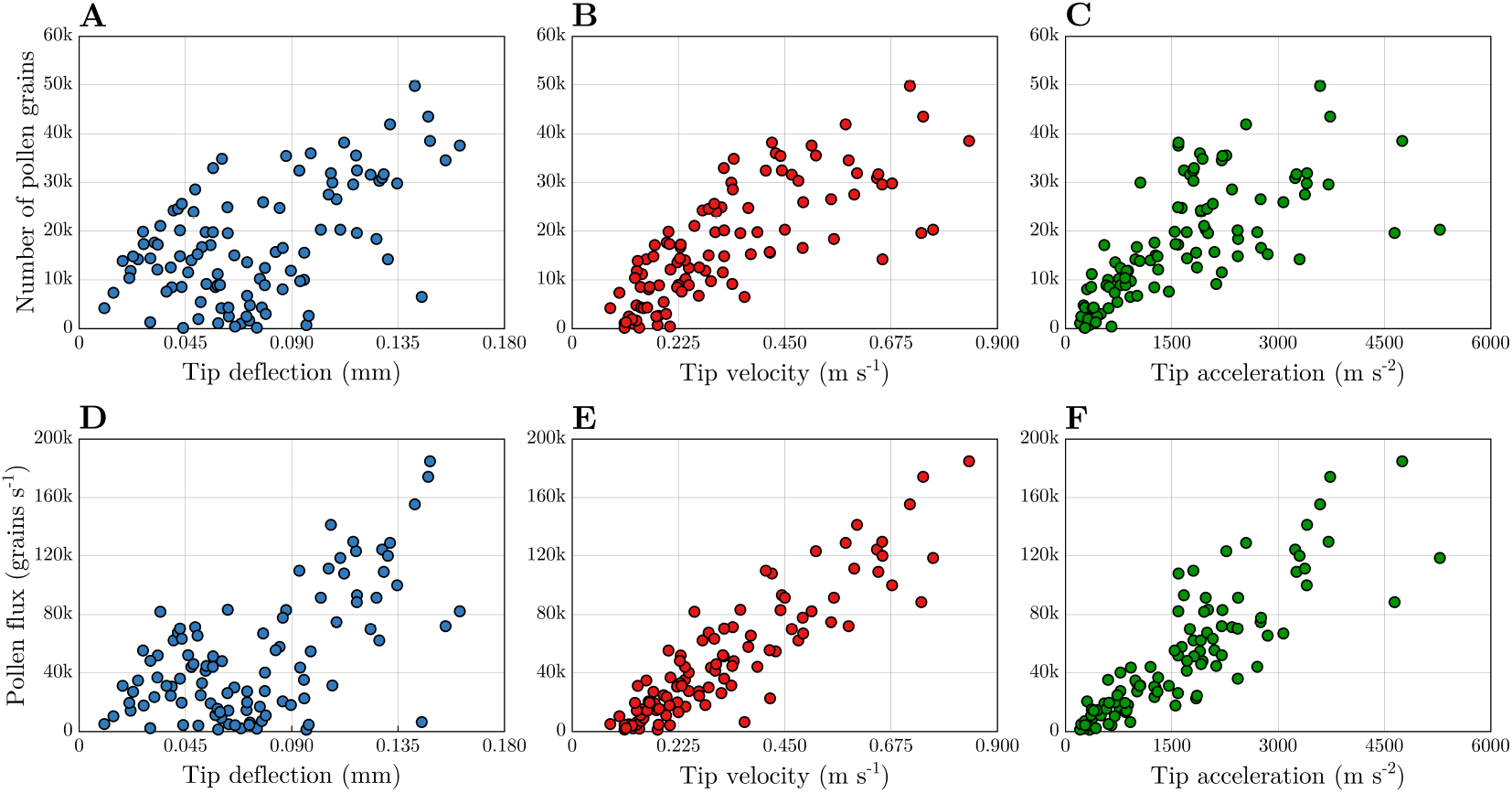
The total number of pollen grains expelled over the 1-second buzzing duration and pollen flux vs. anther tip displacement, velocity and acceleration amplitude. Pollen grains released vs anther tip dynamics (A,B,C). Pollen flux vs anther tip dynamics (D,E,F)

To quantify these relationships, we calculated Pearson correlation coefficients between pollen release metrics and anther tip kinematics. Total pollen released exhibited positive correlations with tip displacement (*r* = 0.516), tip velocity (*r* = 0.755), and tip acceleration (*r* = 0.702), with all correlations significant (*p <* 0.001). Stronger relationships were observed for initial pollen flux, which was correlated with tip displacement (*r* = 0.621), tip velocity (*r* = 0.898), and tip acceleration (*r* = 0.853) (all *p <* 0.001). Among the kinematic measures considered, tip velocity exhibited the strongest correlation with both total pollen released and initial pollen flux.

### 3.4 Pollen Ejection Models

Predictions from the linear and nonlinear pollen release models are compared with the normalized experimental measurements in Fig. 5. For both models, the predicted fraction of pollen released increases nonlinearly with tip velocity and generally defines an upper bound on the experimentally observed pollen release. Consequently, the models tend to overpredict the amount of pollen released for a given tip velocity, particularly at larger velocities. Potential reasons for these overpredictions are considered in the discussion. In contrast, predictions of normalized pollen flux show substantially better agreement with the experimental measurements, with much of the predicted range overlapping the observed data.

**Figure 5:**
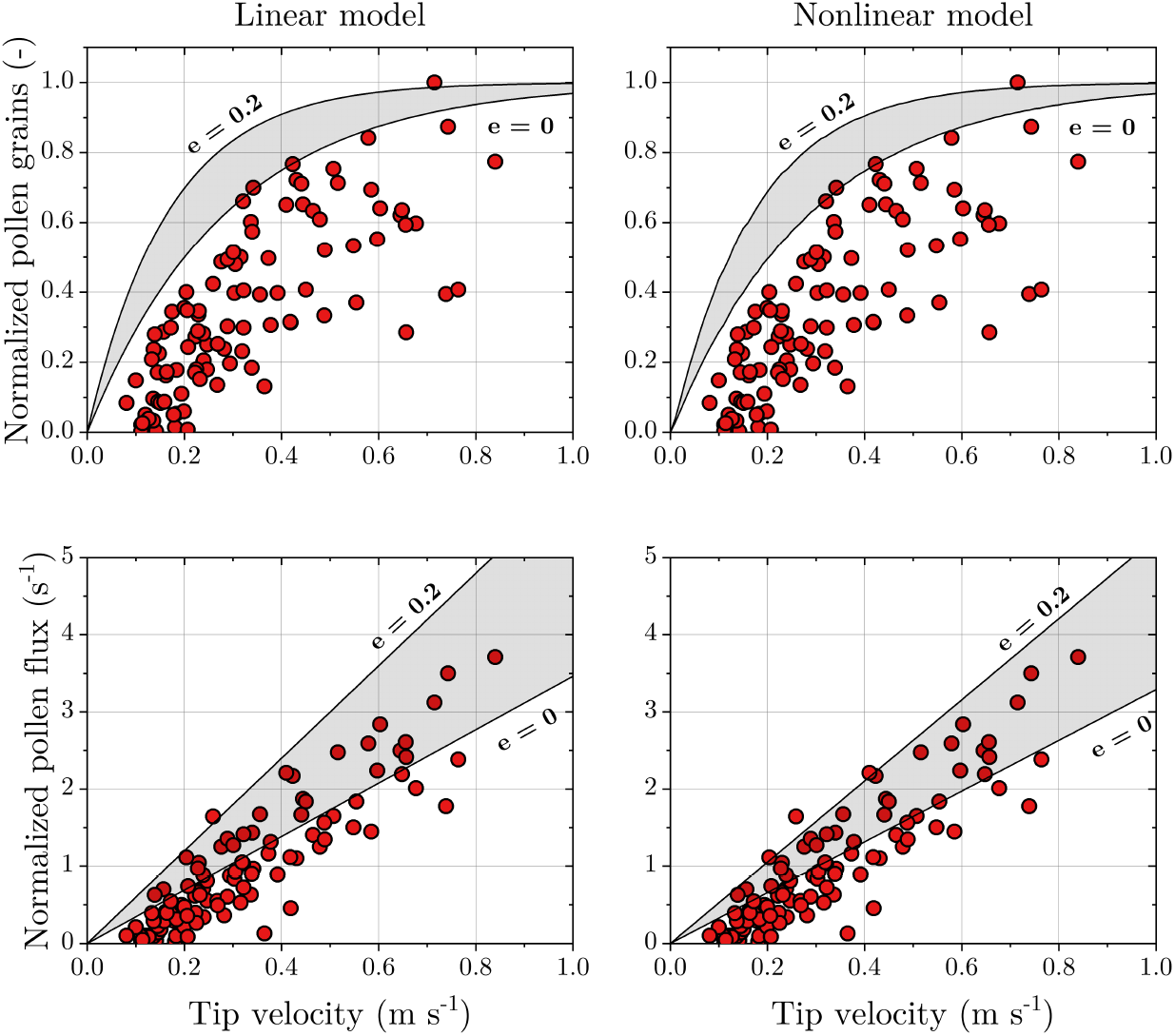
Comparison of experimental pollen release measurements with predictions from the linear and nonlinear statistical pollen release models. Top row: Normalized pollen release as a function of anther tip velocity. Bottom row: Normalized pollen flux as a function of anther tip velocity. Experimental measurements are shown as red circles. Model predictions are shown for coefficients of restitution of *e* = 0 and *e* = 0.2 (solid lines), with shaded regions indicating the range of model predictions bounded by these values. Both models predict increasing pollen release and pollen flux with increasing tip velocity. The linear model closely reproduces the predictions of the nonlinear model while avoiding numerical integration of the governing equations.

The linear and nonlinear models produced similar predictions across the range of tip velocities considered, with the largest discrepancies occurring for the higher coefficient of restitution (*e* = 0.2). This agreement highlights the utility of the analytical approximation, which yields estimates of pollen release metrics without requiring numerical integration of the governing equations. Furthermore, the linear model predicts that pollen flux is directly proportional to anther tip velocity (Eq. 16), consistent with the strong linear relationship observed experimentally.

Despite these successes, both models predict pollen release for any nonzero tip velocity. Experimentally, however, pollen release was not observed below a tip velocity of approximately 100 mm/s, suggesting the existence of a minimum velocity threshold required to initiate pollen motion. Potential explanations for this discrepancy are discussed in the discussion.

## 4 Discussion

This study investigated how excitation frequency influences pollen release from poricidal anthers. Using high-speed videography and particle-tracking methods, we quantified both initial pollen flux and total pollen release during controlled buzzing events. Our results demonstrate that pollen release is strongly influenced by anther structural dynamics, with both pollen release metrics peaking near the anther’s natural frequency. We further show that simple statistical models capture key trends in pollen release but have important limitations. The following sections discuss the implications of these findings for buzz pollination and the utility of statistical models for predicting pollen ejection.

### 4.1 Pollen expulsion depends on excitation frequency

Pollen expulsion, quantified by both the initial pollen flux and the total pollen released during a finite buzzing event, depended strongly on excitation frequency. This finding contrasts with previous studies that concluded there was no optimal buzzing frequency for maximizing pollen release [29, 30, 33]. Here, anther tip displacement, velocity, and acceleration amplitudes were greatest when excitation frequencies approached the anther’s natural frequency, indicating a near-resonant response. Consistent with previous experimental work, total pollen release was most strongly correlated with vibration velocity amplitude rather than displacement or acceleration amplitude [33]. The same trend was observed for initial pollen flux, a quantity that has not previously been reported. These findings suggest that excitation frequency influences pollen release through its effect on anther dynamics, where buzzing near the anther’s natural frequency amplifies vibration velocity and consequently increases both the rate and total amount of pollen released.

Resonance may improve the mechanical efficiency of buzz pollination, which we define as the amount of pollen extracted per unit energy expended by the bee. Near resonance, relatively small forcing amplitudes can generate large structural responses. In the present study, we controlled excitation velocity rather than force and therefore observed increased pollen release when excitation frequencies approached the anther’s natural frequency. However, the same principle can be viewed from the opposite perspective. If anther kinematics were instead held constant across frequencies, the force required to achieve a given vibration amplitude would decrease near resonance. Consequently, lower instantaneous power and less total energy would be required to achieve the same pollen release outcome over a finite buzzing duration. Resonance therefore has the potential to improve pollination efficiency in two complementary ways: by increasing pollen release for a given energetic investment or by reducing the energetic cost required to extract a given amount of pollen.

Presently, there is insufficient evidence that bees target resonance or vibration amplification in natural contexts. The mean natural frequency measured for isolated anthers was approximately 870 Hz, substantially higher than the 100–400 Hz buzzing frequencies typically reported for buzz-pollinating insects [22]. If these values were representative of the *in vivo* bee-flower system, resonance-based amplification would be unlikely to contribute substantially to pollen release. However, several factors suggest that the resonant frequencies experienced during natural pollination may differ from those measured here. First, computational models indicate that pollinator mass can substantially alter anther dynamics. Because the mass of a bee may exceed that of the anther by several orders of magnitude, inclusion of bee mass can reduce the natural frequency of the coupled system [25]. Second, the boundary conditions used in our experiments likely increased structural stiffness. Whereas the anthers were rigidly mounted at their bases, anthers in nature are supported by a compliant filament that may behave as an additional spring element and further reduce natural frequency. Consistent with this interpretation, experimental measurements of entire stamens in other *Solanum* species have reported natural frequencies ranging from 45–300 Hz, substantially lower than the isolated-anther frequencies measured here [42]. Finally, anther turgor pressure also appears to influence natural frequency and is difficult to control experimentally [43]. Because turgor varies throughout the day in response to environmental conditions [44, 45], the natural frequency of an anther may be better viewed as a range of values rather than a fixed quantity.

Taken together, these considerations suggest that vibration amplification remains a plausible mechanism for enhancing pollen extraction during buzz pollination. Importantly, exploiting resonance may not require bees to precisely match a single resonant frequency. Instead, bees may operate within a broad frequency band that overlaps the dynamically shifting resonant range of the flower. Such a strategy would be inherently robust to variation in bee mass, filament compliance, hydration state, and other sources of biological variability. Under this framework, efficient pollen extraction could be achieved through approximate rather than exact frequency matching. This hypothesis is consistent with observations that some buzz-pollinating species adjust their buzzing frequencies in response to floral traits [46]. Rather than tuning to a fixed resonant frequency, bees may modulate their buzzing behavior to remain within a frequency range that promotes vibration amplification and efficient pollen release. Testing this hypothesis will require measurements of the coupled bee-flower system under natural conditions, where interactions among bee mass, floral morphology, and anther hydration can be explicitly quantified. Such studies will help determine the extent to which dynamic coupling influences pollen dispensing and pollination efficiency *in viv*o.

### 4.2 Do simple models accurately model pollen expulsion?

Even relatively simple mathematical models provide a useful framework for understanding the physics governing pollen ejection. Although statistical models lack the fidelity of higher-order approaches based on billiards theory [38] or the discrete element method [40, 41], they permit rapid estimation of pollen release metrics and provide direct insight into the governing mechanisms. By applying simplifying assumptions to the nonlinear statistical model, we derived approximate analytical solutions that identify the key parameters controlling pollen release, including vibration velocity, the ratio of aperture area to interior wall area, and characteristic anther length scales. The resulting analytical model produced predictions that closely matched those of the full nonlinear model while avoiding numerical integration of the governing equations. However, although both statistical models predicted normalized pollen flux with reasonable accuracy, their predictions of total pollen release over a finite buzzing interval were substantially less accurate. These findings suggest that statistical models are best suited for describing the initial stages of pollen ejection, whereas longer-duration predictions may require additional physical mechanisms not captured by the present framework.

Statistical models treat pollen grains as a granular gas and therefore neglect particle-particle interactions such as collisions, aggregation, and clumping. We hypothesize that these interactions become increasingly important as pollen migrates toward the anther aperture during buzzing. Previous discrete element simulations of buzz pollination suggest that particle-particle interactions occur more frequently than particle-wall interactions [40], indicating that collective pollen dynamics may strongly influence pollen transport. In addition, pollen grains are coated with pollenkitt, a viscous substance that may underpin adhesion and clump formation [47, 48]. Electrostatic interactions may also contribute, although their role in buzz-pollinated species remains poorly understood [49]. Such interactions could reduce the effective volume available for particle motion or decrease the effective aperture size through clogging near the anther tip, thereby limiting pollen release. Because statistical models cannot account for these evolving particle-scale effects, they likely overestimate pollen release over extended buzzing durations, consistent with the trends observed in the present study.

Adhesive interactions may also explain why pollen release was not observed at low vibration amplitudes despite being predicted by the statistical models. In the models, pollen grains are assumed to become mobile as soon as energy is transferred from the vibrating anther walls. In reality, however, pollen grains are initially at rest and may adhere to one another or to the interior anther walls through pollenkitt, electrostatic forces, or other adhesive mechanisms. We therefore hypothesize the existence of a critical acceleration threshold that must be exceeded before pollen grains or pollen clumps become mobilized. Physically, this threshold corresponds to the point at which acceleration-induced inertial forces overcome adhesive forces within the pollen mass. Once mobilized, pollen grains can participate in the collision processes described by the statistical models. Because such adhesion-mediated thresholds are not represented in the present framework, the models predict pollen release at arbitrarily small vibration amplitudes and fail to capture the experimentally observed velocity threshold. Higher-fidelity approaches based on the discrete element method are better suited to incorporate these mechanisms, although additional measurements of pollen adhesion properties are needed before such models can be parameterized reliably.

Finally, statistical models are sensitive to the parameters used to populate them and to the experimental measurements against which they are compared. Among the model parameters, the coefficient of restitution is particularly important because it governs the rate at which pollen grains gain and dissipate energy through collisions. However, the coefficient of restitution of pollen grains remains poorly characterized and is likely influenced by factors such as pollen morphology, hydration state, and surface chemistry. Similarly, geometric parameters, including the ratio of aperture area to interior wall area, can vary among species [27]. Additional uncertainty arises from the experimental normalization procedure. Because the initial pollen content of individual anthers was not measured, pollen release was normalized using the maximum number of grains expelled in any trial rather than the true pollen count of each anther. Consequently, the normalized release fractions may not accurately represent the proportion of pollen removed from individual anthers, potentially contributing to variability in the experimental data and discrepancies between model predictions and measurements. Our analytical approximation demonstrates that relatively small changes in these biological and geometric parameters can substantially alter predicted pollen flux and pollen release. Improving the predictive capability of statistical models will require more accurate characterization of pollen collision mechanics, adhesion forces, anther geometry, and initial pollen content.

## Author contributions

Conceptualization – MJ; Data curation – MA; Formal analysis – MA, MJ, BC; Funding Acquisition – MJ; Investigation – MA; Methodology – MA, MJ, SM; Project administration – MJ; Resources – MJ; Software – MA, SM, BC; Supervision – MJ, SM; Validation – MA; Visualization – MA; Writing (original draft) – MA, MJ; Writing (review and editing) – MA, MJ, BC, SM

## Data Availability

The datasets supporting this article have been deposited in Zenodo and are openly available at https://zenodo.org/records/20722407. The repository includes the experimental data compiled into a spreadsheet. Data are available under a Creative Commons Attribution 4.0 International (CC BY 4.0) license.

## References

[1] David J Pritchard and Mario Vallejo-Marín. Buzz pollination. Current Biology, 30(15):R858– R860, 2020.

[2] Mario Vallejo-Marín. Buzz pollination: studying bee vibrations on flowers. New Phytologist, 224(3):1068–1074, 2019.

[3] Stephen L Buchmann and Mark Jankauski. Buzz pollination: Bee bites and floral vibrations. Current Biology, 34(18):R864–R866, 2024.

[4] Mario Vallejo-Marín. How and why do bees buzz? Implications for buzz pollination. Journal of Experimental Botany, pages 1080–1092, 2021.

[5] Carolyn E.B. Proenqa. Buzz pollination-older and more widespread than we think? Journal of Tropical Ecology, 8(1), 1992.

[6] Sophie Cardinal, Stephen L. Buchmann, and Avery L. Russell. The evolution of floral sonication, a pollen foraging behavior used by bees (Anthophila). Evolution, 72(3):590–600, 2018.

[7] Callin M. Switzer, Avery L. Russell, Daniel R. Papaj, Stacey A. Combes, and Robin Hopkins. Sonicating bees demonstrate flexible pollen extraction without instrumental learning. Current Zoology, 65(4):425–436, 2019.

[8] Yuanqing Xu, Bentao Wu, Mario Vallejo-Marin, Petert Bernhardt, Mark Jankauski, Dezhu Li, Stephen Buchmann, Jianing Wu, and Hong Wang. Buzz-pollination leads to size-dependent associations between bumblebees and Pedicularis flowers. ESS Open Archive eprints, 366:36612114, 2024.

[9] S. L. Buchmann. Buzz pollination in angiosperms. In Handbook of Experimental Pollination Biology. 1983.

[10] Paul A. De Luca and Mario Vallejo-Marín. What’s the ‘buzz’ about? The ecology and evolutionary significance of buzz-pollination. Current Opinion in Plant Biology, 16(4):429–435, 2013.

[11] J. R. Matthews and C. M. Maclachlan. The Structure of Certain Poricidal Anthers. Transactions of the Botanical Society of Edinburgh, 30(2), 1929.

[12] Avery L. Russell, Stephen L. Buchmann, and Daniel R. Papaj. How a generalist bee achieves high efficiency of pollen collection on diverse floral resources. Behavioral Ecology, 28(4):991–1003, 2017.

[13] Stephen L. Buchmann and James P. Hurley. A biophysical model for buzz pollination in angiosperms. Journal of Theoretical Biology, 72(4):639–657, 1978.

[14] Marcus J. King and Stephen L. Buchmann. Floral sonication by bees: Mesosomal vibration by Bombus and Xylocopa, but not Apis (Hymenoptera: Apidae), ejects pollen from poricidal anthers. Journal of the Kansas Entomological Society, 76(2):295–305, 2003.

[15] Marcus J. King, Stephen L. Buchmann, and Hayward Spangler. Activity of asynchronous flight muscle from two bee families during sonication (buzzing). Journal of Experimental Biology, 199(10):2317–2321, 1996.

[16] Mario Vallejo-Marin and Avery L. Russell. Harvesting pollen with vibrations: towards an integrative understanding of the proximate and ultimate reasons for buzz pollination. Annals of Botany, 133(3), 2024.

[17] Hazel Cooley and Mario Vallejo-Marín. Buzz-Pollinated Crops: A Global Review and Meta-analysis of the Effects of Supplemental Bee Pollination in Tomato. Journal of economic entomology, 114(2):505–519, 2021.

[18] Avery L. Russell, Stephen L. Buchmann, John S. Ascher, Zhiheng Wang, Ricardo Kriebel, Diana D. Jolles, Michael C. Orr, and Alice C. Hughes. Global patterns and drivers of buzzing bees and poricidal plants. Current Biology, 34(14), 2024.

[19] Blanca Arroyo-Correa, Ceit Beattie, and Mario Vallejo-Marın. Bee and floral traits affect the characteristics of the vibrations experienced by flowers during buzz pollination. Journal of Experimental Biology, 222(4):jeb198176, 2019.

[20] Charlie Woodrow, Noah Jafferis, Yuchen Kang, and Mario Vallejo-Marín. Buzz-pollinating bees deliver thoracic vibrations to flowers through periodic biting. Current Biology, 34(18):4104–4113, 2024.

[21] M. Jankauski, C. Casey, C. Heveran, K. Busby, and S. Buchmann. Carpenter bee thorax vibration and force generation inform pollen release mechanisms during floral buzzing. Scientific Reports, 12(1):12654, 2022.

[22] Paul A. De Luca, Darryl A. Cox, and Mario Vallejo-Marín. Comparison of pollination and defensive buzzes in bumblebees indicates species-specific and context-dependent vibrations. Naturwissenschaften, 101(4):331–338, 2014.

[23] Natacha Rossi, Mario Vallejo-Marin, and Elizabeth Nicholls. First direct quantification of floral handling costs in bees. Proceedings of the Royal Society B: Biological Sciences, 293(2071), 2026.

[24] Emmanuel De Langre. Plant vibrations at all scales: A review. Journal of Experimental Botany, 70(14), 2019.

[25] Mark Jankauski, Riggs Ferguson, Avery Russell, and Stephen Buchmann. Structural dynamics of real and modelled Solanum stamens: implications for pollen ejection by buzzing bees. Journal of The Royal Society Interface, 19(188):20220040, 4 2022.

[26] Mitchell Alvord, Jenna McNally, Cailin Casey, and Mark Jankauski. Turgor pressure affects transverse stiffness and resonant frequencies of buzz-pollinated poricidal anthers. Journal of Experimental Botany, page erae504, 2024.

[27] Jurene E. Kemp and Mario Vallejo-Marín. Pollen dispensing schedules in buzz-pollinated plants: experimental comparison of species with contrasting floral morphologies. American Journal of Botany, 108(6), 2021.

[28] Mario Vallejo-Marín and Anna Lundgren. Gradual pollen release in a buzz-pollinated plant: Investigating pollen presentation theory under bee visitation. Functional Ecology, 2025.

[29] Mandeep Tayal and Rupesh Kariyat. Examining the role of buzzing time and acoustics on pollen extraction of solanum elaeagnifolium. Plants, 10(12), 2021.

[30] Paul A. De Luca, Luc F. Bussière, Daniel Souto-Vilaros, Dave Goulson, Andrew C. Mason, and Mario Vallejo-Marín. Variability in bumblebee pollination buzzes affects the quantity of pollen released from flowers. Oecologia, 172(3):805–816, 2013.

[31] Carlos Eduardo Pereira Nunes and Mario Vallejo-Marín. How much pollen do beelike floral vibrations remove from different types of anthers? International Journal of Plant Sciences, 183(9):768–776, 2022.

[32] Rachel V. Wilkins, Maggie M. Mayberry, Mario Vallejo-Marín, and Avery L. Russell. Hold tight or loosen up? Functional consequences of a shift in anther architecture depend substantially on bee body size. Oecologia, 200(1-2), 2022.

[33] Conrado Augusto Rosi-Denadai, Priscila Cássia Souza Araújo, Lucio Ant ônio de Oliveira Campos, Lirio Cosme, and Raul Narciso Carvalho Guedes. Buzz-pollination in Neotropical bees: genus-dependent frequencies and lack of optimal frequency for pollen release. Insect Science, 27(1):133–143, 2020.

[34] Tanmay Nath, Alexander Mathis, An Chi Chen, Amir Patel, Matthias Bethge, and Mackenzie Weygandt Mathis. Using DeepLabCut for 3D markerless pose estimation across species and behaviors. Nature Protocols, 14(7), 2019.

[35] Johannes Schindelin, Ignacio Arganda-Carreras, Erwin Frise, Verena Kaynig, Mark Longair, Tobias Pietzsch, Stephan Preibisch, Curtis Rueden, Stephan Saalfeld, Benjamin Schmid, Jean Yves Tinevez, Daniel James White, Volker Hartenstein, Kevin Eliceiri, Pavel Tomancak, and Albert Cardona. Fiji: An open-source platform for biological-image analysis, 2012.

[36] Jean Yves Tinevez, Nick Perry, Johannes Schindelin, Genevieve M. Hoopes, Gregory D. Reynolds, Emmanuel Laplantine, Sebastian Y. Bednarek, Spencer L. Shorte, and Kevin W. Eliceiri. Track-Mate: An open and extensible platform for single-particle tracking. Methods, 115, 2017.

[37] Dmitry Ershov, Minh Son Phan, Joanna W. Pylvänäinen, Stéphane U. Rigaud, Laure Le Blanc, Arthur Charles-Orszag, James R.W. Conway, Romain F. Laine, Nathan H. Roy, Daria Bonazzi, Guillaume Duménil, Guillaume Jacquemet, and Jean Yves Tinevez. TrackMate 7: integrating state-of-the-art segmentation algorithms into tracking pipelines. Nature Methods, 19(7), 2022.

[38] Matheus Hansen, Gabriel C. Lanes, Vinícius L.G. Brito, and Edson D. Leonel. Investigation of pollen release by poricidal anthers using mathematical billiards. Physical Review E, 104(3), 2021.

[39] Daniel Borin, Vinicius Lourenço Garcia De Brito, Edson Denis Leonel, and Matheus Hansen. Buzz pollination: A theoretical analysis via scaling invariance. Physical Review E, 110(5), 2024.

[40] Caelen Boucher-Bergstedt, Mark Jankauski, and Erick Johnson. Buzz pollination: investigations of pollen expulsion using the discrete element method. Journal of the Royal Society Interface, 22(222):20240526, 2025.

[41] Qiang Shi, Yong Liu, Bin Wang, Yafei Wang, Xiaoxue Du, Yongzhong Zhang, Hanping Mao, and Xiaoyue Yang. Discrete element simulation of buzz pollination in tomato. Scientific Reports, 15(1):6710, 2025.

[42] Carlos Eduardo Pereira Nunes, Lucy Nevard, Fernando Montealegre-Z, and Mario Vallejo-Marín. Variation in the natural frequency of stamens in six morphologically diverse, buzz-pollinated, heterantherous Solanum taxa and its relationship to bee vibrations . Botanical Journal of the Linnean Society, pages 541–553, 2021.

[43] Benjamin S. Lazarus and Agnes S. Dellinger. Wilting may leave bees wanting: drops in turgor pressure may reduce viability of buzz-pollinated flowers, 2025.

[44] Wieland Fricke. Cell turgor, osmotic pressure and water potential in the upper epidermis of barley leaves in relation to cell location and in response to NaCl and air humidity. Journal of Experimental Botany, 48(306):45–58, 1997.

[45] James O Eubanks. Effect of light intensity and osmotic stress on the water relations of Populus tremuloides. Forest Science, 17(1):79–82, 1971.

[46] Sarah A. Corbet and Shuang Quan Huang. Buzz pollination in eight bumblebee-pollinated Pedicularis species: Does it involve vibration-induced triboelectric charging of pollen grains? Annals of Botany, 114(8):1665–1674, 2014.

[47] Shuto Ito and Stanislav N. Gorb. Attachment-based mechanisms underlying capture and release of pollen grains. Journal of the Royal Society Interface, 16(157), 2019.

[48] Haisheng Lin, Ismael Gomez, and J. Carson Meredith. Pollenkitt wetting mechanism enables species-specific tunable pollen adhesion. Langmuir, 29(9), 2013.

[49] Sarah A. Corbet, James Beament, and D Eisikowitch. Are electrostatic forces involved in pollen transfer? Plant, Cell & Environment, 5(2), 1982.

